# *DYRK1A*-related intellectual disability: a syndrome associated with congenital anomalies of the kidney and urinary tract

**DOI:** 10.1101/516856

**Authors:** Alexandria T.M. Blackburn, Nasim Bekheirnia, Vanessa C. Uma, Jill A. Rosenfeld, Matthew N. Bainbridge, Yaping Yang, Pengfei Liu, Suneeta Madan-Khetarpal, Mauricio R. Delgado, Louanne Hudgins, Ian Krantz, David Rodriguez-Buritica, Patricia G. Wheeler, Lihadh Al Gazali, Aisha Mohamed Saeed Mohamed Al Shamsi, Natalia Gomez-Ospina, Hsiao-Tuan Chao, Ghayda M. Mirzaa, Angela E. Scheuerle, Mary K Kukolich, Fernando Scaglia, Christine Eng, Michael C. Braun, Dolores J. Lamb, Rachel K. Miller, Mir Reza Bekheirnia

**Affiliations:** Department of Pediatrics, Pediatric Research Center, University of Texas Health Science Center, McGovern Medical School, Houston, TX 77030; Program in Genetics and Epigenetics, The University of Texas MD Anderson Cancer Center University of Texas Health Science Center Graduate School of Biomedical Sciences, Houston, TX 77030; Renal Section, Department of Pediatrics, Baylor College of Medicine, Houston, TX, 77030; Texas Children’s Hospital, Houston, TX 77030; Baylor College of Medicine, Houston, TX 77030; Department of Molecular and Human Genetics, Baylor College of Medicine, Houston, TX 77030; Codified Genomics, LLC, Houston, TX 77030; Rady Children’s Institute for Genomic Medicine, San Diego, CA, 92123; Baylor Genetics, Houston, Texas 77030; Department of Pediatrics, University of Pittsburgh School of Medicine, Pittsburgh, PA 15224; Department of neurology, The University of Texas Southwestern Medical Center, Dallas, TX 75390; Department of Pediatrics, Division of Medical Genetics, Stanford University, Stanford, California 94305; Department of Pediatrics, Perelman School of Medicine at the University of Pennsylvania, Philadelphia, PA 19104; Department of Pediatrics, McGovern Medical School, The University of Texas Health Science Center at Houston, TX 77030; Arnold Palmer Hospital, Orlando, FL.; Department of Paediatrics, College of Medicine & Health Sciences, UAE university, UAE; Paediatrics Department, Tawam Hospital, Al-Ain, United Arab Emirates; Department of Pediatrics (Child Neurology), Baylor College of Medicine, Houston, TX 77030; Center for Integrative Brain Research, Seattle Children’s Research Institute, Seattle, Washington, USA; Department of Pediatrics, University of Washington, Seattle, Washington, USA; Department of Pediatrics (Genetics and Metabolism), The University of Texas Southwestern Medical Center, Dallas, TX 75390; Clinical Genetics, Cook Children’s Medical Center, Fort Worth, TX 76104; BCM-CUHK Center of Medical Genetics, Prince of Wales Hospital, ShaTin, Hong Kong SAR; Department of Urology and Center for Reproductive Genomics, Weill Cornell Medicine, New York, NY 10065; Department of Genetics, University of Texas MD Anderson Cancer Center, Houston, Texas 77030; Program in Biochemistry and Cell Biology, The University of Texas MD Anderson Cancer Center University of Texas Health Science Center Graduate School of Biomedical Sciences, Houston, Texas 77030

**Keywords:** CAKUT, exome sequencing, *DYRK1A*, *Xenopus*, intellectual disability

## Abstract

**Purpose:** Haploinsufficiency of *DYRK1A* causes a recognizable clinical syndrome. The goal of this paper is to investigate congenital anomalies of the kidney and urinary tract (CAKUT) and genital defects (GD) in patients with *DYRK1A* mutations.

**Methods:** A large database of clinical exome sequencing (ES) was queried for *de novo DYRK1A* mutations and CAKUT/GD phenotypes were characterized. *Xenopus laevis* (frog) was chosen as a model organism to assess Dyrk1a’s role in renal development.

**Results:** Phenotypic details and mutations of 19 patients were compiled after an initial observation that one patient with a *de novo* pathogenic mutation in *DYRK1A* had GD. CAKUT/GD data were available from 15 patients, 11 of whom present with CAKUT/GD. Studies in *Xenopus* embryos demonstrate that knockdown of Dyrk1a disrupts the development of segments of developing embryonic nephrons, which ultimately give rise to the entire genitourinary (GU) tract. These defects could be rescued by co-injecting wildtype human *DYRK1A* RNA, but not with truncated *DYRK1A*^*R205*\*^ RNA.

**Conclusion:** Evidence supports routine GU screening of all individuals with *de novo DYRK1A* pathogenic variants to ensure optimized clinical management. Collectively, the reported clinical data and loss of function studies in *Xenopus* substantiate a novel role for *DYRK1A* in GU development.

## INTRODUCTION

The dual-specificity tyrosine-phosphorylation-regulated kinase (DYRK) family of protein kinases are conserved across species from lower eukaryotes to mammals.^1^ DYRK family members are activated by autophosphorylating a tyrosine residue in their activation loop.^2^ DYRK1A is the most extensively characterized member of the DYRK family, which in humans, is encoded by the *DYRK1A* gene located in the Down syndrome critical region of chromosome 21.^3^ A growing body of literature implicates a strong causal relationship between *DYRK1A* haploinsufficiency and a recognizable syndrome known as *DYRK1A*-related intellectual disability syndrome.^3^–^5^ In addition to intellectual disability (ID), other frequently occurring features include; intrauterine growth retardation (IUGR), difficulty feeding (DF) with failure to thrive (FTT), microcephaly, seizures, dysmorphic facial features, and developmental delays (DD),^5^ while additional phenotypic features are observed less commonly.^5,6^ While many features of this syndrome are well-characterized; the full phenotypic spectrum has yet to be defined. This manuscript presents a cohort of individuals with *de novo* (when both parental samples available) *DYRK1A* single nucleotide variants (SNVs) or small deletions ≤10 base pairs and defines CAKUT/GD that have not been previously described in patients with *DYRK1A* syndrome. We also provide supporting evidence, using *Xenopus laevis* embryos as a model, that *DYRK1A* is required for GU development and that a pathogenic variant of human *DYRK1A* is directly responsible for the CAKUT/GD phenotype. Together, these findings support the investigation of potential CAKUT/GD in the clinical workup of patients with *DYRK1A*-related ID syndrome.

## MATERIALS AND METHODS

### Study participants

The index patient was seen in the Renal Genetics Clinic (RGC) at Texas Children’s Hospital (TCH). Subsequently, patients who had ES in a clinical diagnostic laboratory [Baylor Genetics (BG)] were queried for *de novo* (except P1, P12 and P15) pathogenic [except P10 (likely pathogenic), P5, P7, and P17 called as VUS in the initial report] variants in *DYRK1A*. Inclusion criteria also included: 1) variant confirmation using Sanger sequencing, and 2) lack of other mutations that could explain the phenotype observed. Exclusion criteria included multiple additional candidate genes that may be related to the phenotype. The final size of our cohort after applying those filters is 19 patients. Clinical phenotype information was collected from initial ES requisition form or contacting referring physicians. The Institutional Review Board approved the study protocol for the Protection of Human Subjects at Baylor College of Medicine.

### ES and data analysis

ES was performed by previously published methods at BG.^7,8,9^ In brief, an Illumina paired-end pre-capture library was constructed with 1ug of DNA, according to the manufacturer’s protocol (Illumina Multiplexing_SamplePrep_Guide_-1005361_D), with modifications as described in the BCM-HGSC Illumina Barcoded Paired-End Capture Library Preparation protocol.^7^ Four pre-captured libraries were pooled and then hybridized in solution to the HGSC CORE design (52Mb, NimbleGen) according to the manufacturer’s protocol NimbleGen SeqCap EZ Exome Library SR User’s Guide (Version 2.2), with minor revisions. Sequencing was performed in paired-end mode with the Illumina HiSeq 2000 platform, with sequencing-by-synthesis reactions extended for 101 cycles from each end with an additional cycle for the index read. With a sequencing yield of 12 Gb, 92% of the targeted exome bases were covered to a depth of 20X or greater. Illumina sequence analysis was performed with the HGSC Mercury analysis pipeline (https://www.hgsc.bcm.edu-/software/mercury)which moves data through various analysis tools from the initial sequence generation on the instrument to annotated variant calls (SNVs and intra-read indels). Variant interpretation was performed according to the most recent guidelines published by the American College of Medical Genetics and Genomics (ACMG).^10^ Accordingly, only variants that met strict criteria were called pathogenic. Sanger sequencing confirmed all variants reported in this paper.

### *Xenopus laevis* embryos and microinjections

*Xenopus* females were induced to produce eggs by injection of human chorionic gonadotropin. Eggs were obtained by standard means, placed in 0.3x MMR (Marc’s modified Ringers solution, 33 mM NaCl, 0.66 mM KCl, 0.33 mM MgSO4, 0.66 mM CaCl2, 1.66 mM HEPES pH 7.4) and fertilized *in vitro*. Blastula cleavage stages and dorsal versus ventral polarity were determined by established methods.^11^ Microinjections were targeted to the V2 blastomere at the eight-cell stage, which provides major contributions to the development of the pronephros.^12^–^14^ Ten nL of injection mix (described below) was injected into embryos. Ten ng of Dyrk1a morpholino 5’-TGCATCGTCC-TCTTTCAAGTCTCAT-3′^15^ or Standard morpholino 5′-CCTCTTACCTCAGTTACAATTTATA-3′ was co-injected with 50 pg RNA (either control *β-galactosidase*, human *DYRK1A*, or patient mutant *DYRK1A* ^*R205\**^) along with 1 ng membrane-RFP RNA^16^ as a lineage tracer to verify that the correct blastomere was injected.

### Immunostaining

Embryos were staged,^11^ fixed, and immunostained^17^ using established protocols. Proximal tubule lumens were labeled with the antibody 3G8 (1:30, European *Xenopus* Resource Centre, Portsmouth, United Kingdom), while the cell membranes of distal and connecting tubules were labeled with antibody the 4A6 (1:5, European *Xenopus* Resource Centre, Portsmouth, United Kingdom).^18^ Rabbit anti-red fluorescent protein (anti-RFP) (1:250, MBL International, Woburn, MA, USA) antibody was used to detect RFP expression of the lineage tracer. Goat anti-mouse or anti-rabbit conjugated to Alexa Fluor 488 or Alexa Fluor 555 (1:500, Invitrogen, Carlsbad, CA, USA) secondary antibodies were used to visualize antibody staining.

### Construct design and *in vitro* transcription

Cloning for Western blot analysis was executed by generating PCR cDNA from a pCS2-*Xenopus*-Dyrk1a-HA construct^15^ using a Bio-Rad C1000 Touch™ Thermal Cycler (Bio-Rad, Hercules, CA, USA), following Addgene’s protocol for PCR cloning. PCR with OneTaq (NEB, Ipswich, MA, USA) was used to amplify wildtype *Xenopus dyrk1a* excluding the nucleotides coding for the stop codon. Forward primer 5’-TTTGGATCCATGCATACAGGAGGAGAGAC-3’ was used to create the control *dyrk1a* construct while forward primer 5’-TTTGGATCCATGA-GACTTGAAAGAGGACGATGCATACAGGAGGAGAGAC-3’ was used to create the *dyrk1a* + 5’ UTR construct, which added part of Dyrk1a’s endogenous 5’ UTR that is recognized by the Dyrk1a MO. The same reverse primer 5’-TTTTTTCTAGACGAGCTTGCCACAGGACTCTG-3’ was used to generate both cDNA fragments. BamHI and XbaI were used to insert cDNA fragments into a pCS2-GFP vector, placing the construct immediately upstream and in-frame with a GFP tag. Cloning for the rescue experiment was executed by traditional cloning methods and PCR cloning. The wildtype human *DYRK1A* cDNA was cut from the pMH-SFB-DYRK1A construct (Addgene, Cambridge, MA, USA) and inserted into pCS2-HA vector using XhoI and contains a gateway vector site 5’ to the cDNA. PCR cloning was used to generate the *DYRK1A*^*R205\**^ truncated mutation. Forward primer 5’-AATGCGGAATTCAGACAAG-TTTGTACAAAAAAGCAGGC-3’ and reverse primer 5’-TAGAGGCTCGAGTCACACTTCTATC-TGTGCTTGATTCAG-3’ were used to amplify the truncated cDNA plus the gateway vector site 5’ to the cDNA from the wildtype human pCS2-HA-DYRK1A vector. The truncated cDNA was then inserted into a pCS2-HA vector using EcoRI and XhoI sites. Prior to *in vitro* transcription, the reading frame of each construct was confirmed by Sanger sequencing (Eurofins, Luxembourg). The cDNA templates were linearized for transcription using NotI. Capped RNAs encoding the constructs were generated using the SP6 mMessage mMachine Kit (Thermo Fisher Scientific, Waltham, MA, USA). The transcribed RNA products were evaluated by a 1% agarose gel using RNA gel loading dye and by optical density (OD 280/260) obtained from a NanoDrop® ND-1000 Spectrophotometer (Thermo Fisher Scientific, Waltham, MA, USA).

### Western blots

To validate knockdown of protein expression by Dyrk1a MO, Western blot analysis was carried out on lysates derived from embryos injected at the single cell stage. Forty ng of either Dyrk1a morpholino 5-TGCATCGTCCTCTTTCAAGTCTCAT-3′^15^ or standard morpholino 5′-CCTCTTACCTCAGTTACAATTTATA-3′ (Gene Tools LLC, Philomath, OR, USA) was co-injected with 1 ng RNA (either *Xenopus dyrk1a* control, or *Xenopus dyrk1a* + 5’ UTR) along with 1 ng membrane-RFP RNA.^16^ Ten-twenty embryos were collected at neurula stages 15-20^11^ to make protein lysates as previously described.^19^ One embryo equivalent of lysate was run in each well of a 7.5% SDS-PAGE gel. Protein was transferred onto a 0.45 µm PVDF membrane (Thermo Fisher Scientific, Waltham, MA, USA) and blocked overnight in KPL block (SeraCare, Milford, MA, USA) at 4°C. Blots were incubated in rabbit anti-green fluorescent protein (anti-GFP) (1:500, iclLab, Portland, OR, USA) for 24 hours at 4°C or rabbit anti glyceraldehyde 3-phosphate dehydrogenase (anti-GAPDH) (1:1000, Santa Cruz, Dallas, TX, USA) for 3 hours at room temperature. Blots were washed with TBS-T, incubated in goat anti-rabbit IgG horseradish peroxidase secondary antibody (1:3000; Bio-Rad, Hercules, CA, USA) for 1 hour at room temperature, and washed again with TBS-T prior to imaging with Bio-Rad ChemiDoc XRS+(Bio-Rad, Hercules, CA, USA) using SuperSignal West Pico PLUS Chemiluminescent Substrate (Thermo Fisher Scientific, Waltham, MA, USA).

### Imaging

Embryos were scored and photographed using an Olympus SZX16 fluorescent stereomicroscope and Olympus DP71 camera (Olympus, Toyko, Japan). 3G8/4A6 immunostained kidney images were taken using a Zeiss LSM800 confocal microscope (Zeiss, Oberkochen, Germany). Fixed embryos were cleared in a solution of BABB/Murray’s clearing solution for confocal imaging (1:2 volume of benzyl alcohol to benzyl benzoate). Images were processed with Adobe Photoshop.

## RESULTS

### CAKUT/GD identified in patients with *DYRK1A* mutations

The index patient was seen in Renal Genetics Clinic (RGC) for the evaluation of ID, global DD, hypospadias, and congenital chordee. Trio ES (tES) revealed a novel *de novo* pathogenic p.G168fs single base pair deletion in *DYRK1A*. A subsequent query of the ES database at BG revealed a total of 18 additional individuals with SNVs or deletions ≤10 base pairs (as defined in methods) in *DYRK1A* among approximately 8000 probands. Phenotype and molecular information of these patients are summarized in Table 1 and Figure 1. Probands were mostly children ranging from 2 to 27 years of age. All 19 of these individuals had neurodevelopmental phenotypes consistent with loss-of-function of *DYRK1A* [MIM: 614104]. We subsequently contacted all referring physicians to obtain further details regarding the CAKUT/GD phenotypes. However, CAKUT/GD statuses of four patients remain unknown. Eleven out of fifteen (73%) individuals with available information presented with CAKUT including unilateral renal agenesis (URA), and GD included undescended testis, hypospadias, etc. (Table 2). One patient (P6) with URA was identified after this newly acquired association of *DYRK1A* with CAKUT was discussed with the referring physician.^20^ Probability of loss-of-function intolerance (pLI) score of *DYRK1A* is 1, indicating that this gene is intolerant to loss-of-function variants.^21^

**Table 1.** Demographics, molecular data, and phenotype of 19 patients with SNVs in DYRK1A identified by clinical exome sequencing (ES)

| Patient number | Age (years) | Gender | Exon number | Nucleotide change | AA change | Novel Variant | DF/FTT | FD | Microcephaly | Seizures | ID | DD | Renal/GU | ASD |
| --- | --- | --- | --- | --- | --- | --- | --- | --- | --- | --- | --- | --- | --- | --- |
| P1* | 2.3 | F | Exon 4 | c.452dupT | p.N151fs | - | + | + | + | - | ukn | M,S | + | ukn |
| P2* | 3.3 | M | Exon 4 | c.461delA | p.K154fs | - | + | + | + | + | + | M,S | + | + |
| P3 | 9.5 | F | Exon 4 | c.489_495del | p.L164fs | + | + | + | + | + | + | M,S | + | + |
| P4 | 11 | M | Exon 4 | c.501delA | p.G168fs | + | - | + | + | + | + | G | + | ukn |
| P5 | 2 | M | Exon 5 | c.517G>T | p.V173F | + | + | + | + | + | ukn | S | nRUS | ukn |
| P6* | 5.1 | M | Exon 5 | c.613C>T | p.R205X | - | - | + | + | - | ukn | U | + | + |
| P7* | 7 | F | Exon 6 | c.734T>G | p.L245R | - | - | + | + | - | + | M,S | + | ukn |
| P8 | 20.5 | M | Exon 6 | c.787C>T | p.R263X | - | + | + | + | + | + | M,S | + | + |
| P9 | 5.7 | M | Exon 7 | c.986_995del | p.S329fs | + | + | + | + | + | + | G | + | + |
| P10 | 9.8 | M | Exon 7 | c.1042G>A | p.G348R | + | - | + | ukn | + | + | G | nRUS | ukn |
| P11 | 13.5 | M | Intron 7 | c.1098+1G>A | N/A | - | - | + | + | + | ukn | G | ukn | + |
| P12 | 27.1 | M | Exon 8 | c.1162dupG | p.A388fs | - | + | ukn | + | + | + | G | + | ukn |
| P13 | 14.8 | F | Exon 8 | c.1217_1220del | p.K406fs | + | + | ukn | ukn | + | ukn | M | ukn | + |
| P14 | 13.8 | F | Exon 9 | c.1309C>T | p.R437X | - | - | ukn | + | - | + | M,S | ukn | ukn |
| P15 | 10.1 | F | Exon 9 | c.1309C>T | p.R437X | - | - | + | + | + | ukn | S | nRUS | ukn |
| P16 | 5.7 | F | Exon 9 | c.1309C>T | p.R437X | - | - | + | + | + | + | G | nRUS | ukn |
| P17 | 19.6 | F | Exon 9 | c.1400G>A | p.R467Q | - | + | + | + | - | + | G | ukn | ukn |
| P18* | 18.5 | M | Exon 9 | c.1399C>T | p.R467X | - | - | + | + | + | + | M,S | + | + |
| P19 | 9.4 | M | Exon 9 | c.1478dupT | p.S494fs | + | ukn | + | + | + | + | M,S | + | + |

ASD: Autism spectrum disease, DD: Developmental delays; Global (G), Motor (M), Speech (S), Unspecified (U), DF/FTT: Difficulty feeding / failure to thrive, FD: Facial dysmorphism, GU: Genitourinary; Normal renal ultrasound nRUS, Unknown ukn, + denotes phenotype observed (see Table 2 for more details), ID: Intellectual disability
\*denotes published patients

**Figure 1.**
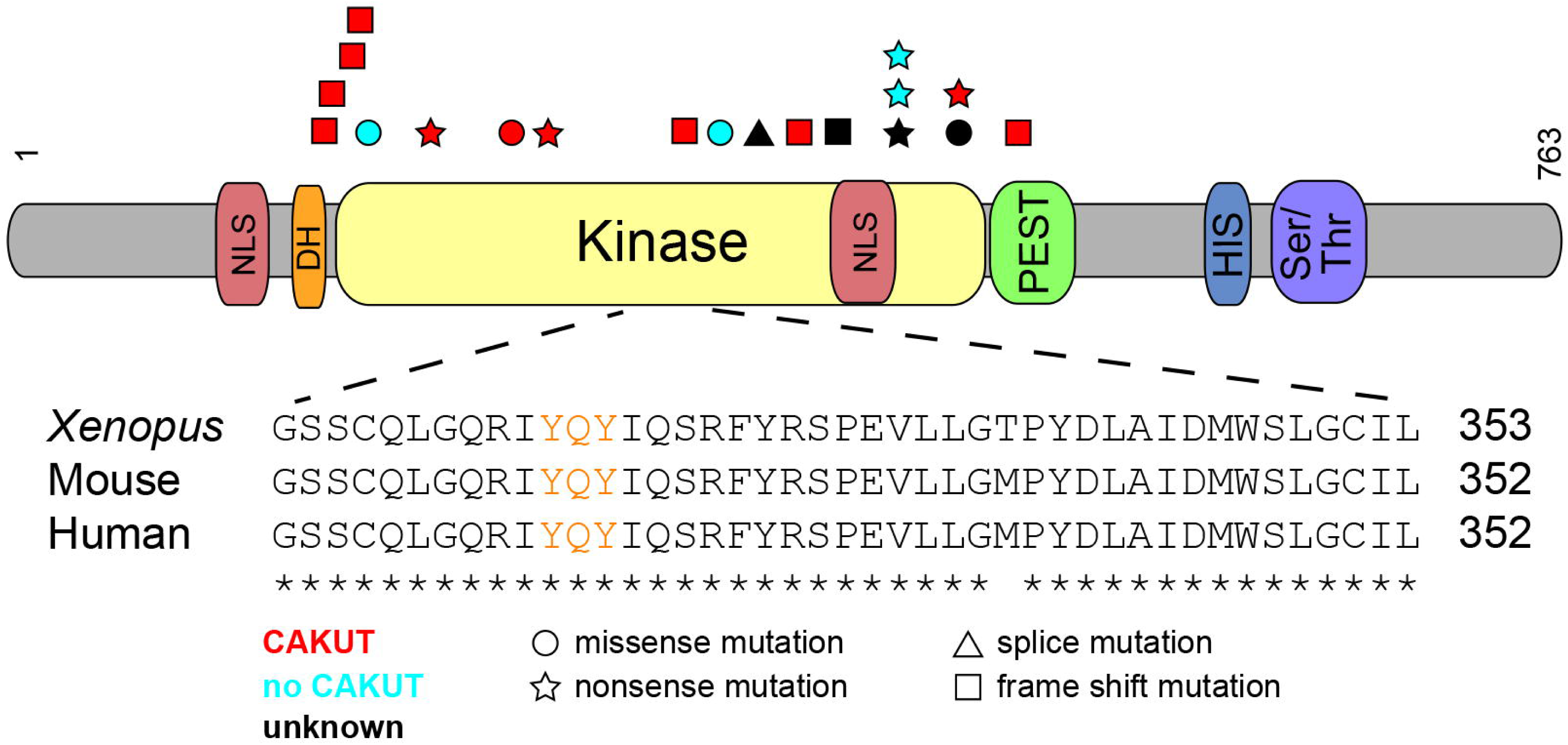
CAKUT associated with DYRK1A mutations in patients with DYRK1A-related intellectual disability syndrome. Schematic shows the DYRK1A protein domains. Shapes, which identify the type of mutation (squares=frame shift mutations, circles=missense mutations, stars=nonsense mutations, triangles=splice mutations), are positioned where *DYRK1A* patient mutations impact the amino acid sequence. Patient mutations are listed in the order they appear in Tables 1 and 2. Mutations that result in CAKUT are red, those that do not result in CAKUT are blue, and those in which the effects on CAKUT status are unknown are black. Inset shows highly conserved sequence surrounding the activation loop (labeled in orange) of the kinase domain.

**Table 2.** Available information about genitourinary (GU) phenotype of patients reported in this study. Eleven patients have GU phenotype. This strongly suggests an important role for DYRK1A in GU tract development.

| Case Number | SNV | Segregation | Renal or GU Phenotype |
| --- | --- | --- | --- |
| P1 | p.N151fs | Mother negative, father's sample unavailable | **Mild unilateral pelviectasis and frequent UTIs** |
| P2 | p.K154fs | *De Novo* | **Genital anomalies** |
| P3 | p.L164fs | *De Novo* | **Kidney abnormalities (not specified)** |
| P4 | p.G168fs | *De Novo* | **Hypospadias, micropenis, and congenital chordee** |
| P5 | p.V173F | *De Novo* | Renal ultrasound is normal with normal genitalia on exam |
| P6 | p.R205X | *De Novo* | **Left renal agenesis** |
| P7 | p.L245R | *De Novo* | **Left renal agenesis** |
| P8 | p.R263X | *De Novo* | **Shawl scrotum and history bilateral orchiopexy** |
| P9 | p.S329fs | *De Novo* | **Hypospadias and kidney abnormalities (tiny echogenic foci)** |
| P10 | p.G348R | *De Novo* | Normal renal ultrasound |
| P11 | c.1098+1G>A | *De Novo* | Unknown |
| P12 | p.A388fs | Mother negative, Father is mosaic | **Frequent UTI** |
| P13 | p.K406fs | *De Novo* | Unknown |
| P14 | p.R437X | *De Novo* | Unknown |
| P15 | p.R437X | Mother negative, father's sample unavailable | Normal renal ultrasound |
| P16 | p.R437X | *De Novo* | Normal renal ultrasound |
| P17 | p.R467Q | *De Novo* | Unknown |
| P18 | p.R467X | *De Novo* | **Orchiopexy and inguinal hernia** |
| P19 | p.S494fs | *De Novo* | **Bilateral inguinal hernias but no renal ultrasound** |

### A majority of the variants found in *DYRK1A* are loss-of-function and are found in the kinase domain

Many of the variants found in this cohort are found in the kinase domain [14 out of 17 (82%)], which spans from residues 159-479, and 11 out of 17 (65%) are thought to undergo nonsense-mediated decay (NMD), as they result in premature stop codons (https://nmdpredictions.shinyapps.io/shiny/) (Figure 1). Of the remaining six variants, one affects splicing, one escapes NMD (p.S494fs), and four are missense variants. The four missense variants are all found in the kinase domain in four individuals. Of these four individuals with missense variants, P7 was diagnosed with URA (p.L245R), P5 and P10 had normal renal ultrasounds (p.V173F and p.G348R), and P18 has an unknown CAKUT status (R467Q).

Because these variants still lead to other *DYRK1A*-syndrome features such as ID they may be important for the catalytic activity or conformational stability of DYRK1A. In fact, a patient in a separate DYRK1A structural study had the same R467Q variant, and this residue was found to be a part of a network of electrostatic interactions thought to play a role in the stability of the DYRK1A protein.^22^ Out of the three mutations not found in the kinase domain, two are just N-terminal to it (N151fs, K154fs) and the last is found just C-terminal to it in the PEST domain (S494fs). Though, theoretically, all mutations that are more N-terminal should result in NMD, a majority of the mutations reside in the kinase domain for unknown reasons that should be studied further.

### *Xenopus laevis* as a model of GU development

DYRK1A’s amino acid sequence is highly conserved among amniotes (https://www.ncbi.nlm.nih.gov/homologene). Even though the N- and C-terminal regions diverge in invertebrates, the amino acid sequence of the kinase domains are similar indicating the importance of this protein throughout evolution. In order to model CAKUT/GD associated with human *DYRK1A* loss-of-function mutations using *Xenopus laevis* embryos (hereafter referred to simply as *Xenopus*), we analyzed the conservation of the whole DYRK1A protein sequence, focusing on the kinase domain. Human and *Xenopus* DYRK1A proteins are 91.3% identical over the entire amino acid sequence (Figure S1). Additionally, the kinase domain, which is where a majority of the mutations that cause CAKUT/GD (in this study) are found, is 97.5% identical (Figure 1). Importantly, the human and *Xenopus* kinase activation loop sequences, which is essential for the kinase activity of DYRK1A, are identical.

*Xenopus* produces large clutch sizes with hundreds of embryos that develop externally, and unilateral embryo injections allow for tissue-targeted knockdowns that are specific to organs such as the kidney.^12^ Their embryonic kidney, the pronephros, can easily be visualized and imaged through a transparent epidermis, and they develop a fully functional kidney in ∼56 hours.^23^ *Xenopus* was chosen because it is an established model of nephron development, and gene expression studies demonstrate that the developing *Xenopus* nephron is anatomically and functionally similar to the mammalian nephron.^24,25^ The embryonic pronephros is the precursor to the mesonephric and metanephric kidney in mammals, and subsequent GU development is dependent upon this structure. Specifically, as the pronephros extends toward the cloaca, the mesonephric nephrons form adjacent to the elongating nephric duct, also known as the Wolffian duct.^26^ The ureteric bud, which is required for the development of the collecting duct system in mammals, then branches from this duct. Because the Wolffian duct is required for GU development in males and Müllerian duct elongation, necessary for normal female anatomy, depends upon the development of the Wolffian duct,^27,28^ the development of the pronephros is critical to both renal and genital development in mammals. Thus, although it is not a well-established model for studying genital formation, the *Xenopus* pronephros is essential for development of the mesonephric/Wolffian duct and subsequently the Müllerian duct, which are required for GU development.^29,30^

### *In vivo* knockdown studies in *Xenopus* assess the role for *dyrk1a* in renal development

To examine *dyrk1a*’s role in renal development in *Xenopus*, anti-sense morpholinos (MO) that block translation were used to knock down endogenous Dyrk1a protein expression. First, the MO was validated by demonstrating through Western blot analysis using a GFP antibody that it correctly targets *Xenopus dyrk1a* RNA. Two constructs were generated with a GFP tag to demonstrate that the MO targets the endogenous 5’ untranslated region (UTR) of *dyrk1a’s* transcript. One construct contains the endogenous transcript’s 5’ UTR (Figure 2A, bottom), while the other construct lacks the 5’ UTR and serves as a positive control (Figure 2A, top). Western blot analysis confirmed that reduction of exogenous Dyrk1a protein levels occurs in neurula stage embryos injected with Dyrk1a MO and *dyrk1a* + 5’ UTR RNA (Figure 2B, lane 5) compared to the Standard MO and *dyrk1a* + 5’ UTR (Figure 2B, lane 4) and Dyrk1a MO or Standard MO plus control RNA (Figure 2B, lane 3, lane 2). Next, knockdown experiments were used to determine whether loss of Dyrk1a function in *Xenopus* results in disruption of kidney development. Embryos were injected into a single V2 blastomere to target a single kidney, leaving the other as an internal control. Knockdown with a Dyrk1a MO resulted in abnormal pronephroi when immunostained with antibodies 3G8 and 4A6, which label the proximal tubules and the distal and connecting tubules (nephric duct), respectively (Figure 3B and B’). Loss of Dyrk1a primarily affected the proximal and distal tubules, with defects seen less frequently in the connecting tubules (nephric duct).

**Figure 2.**
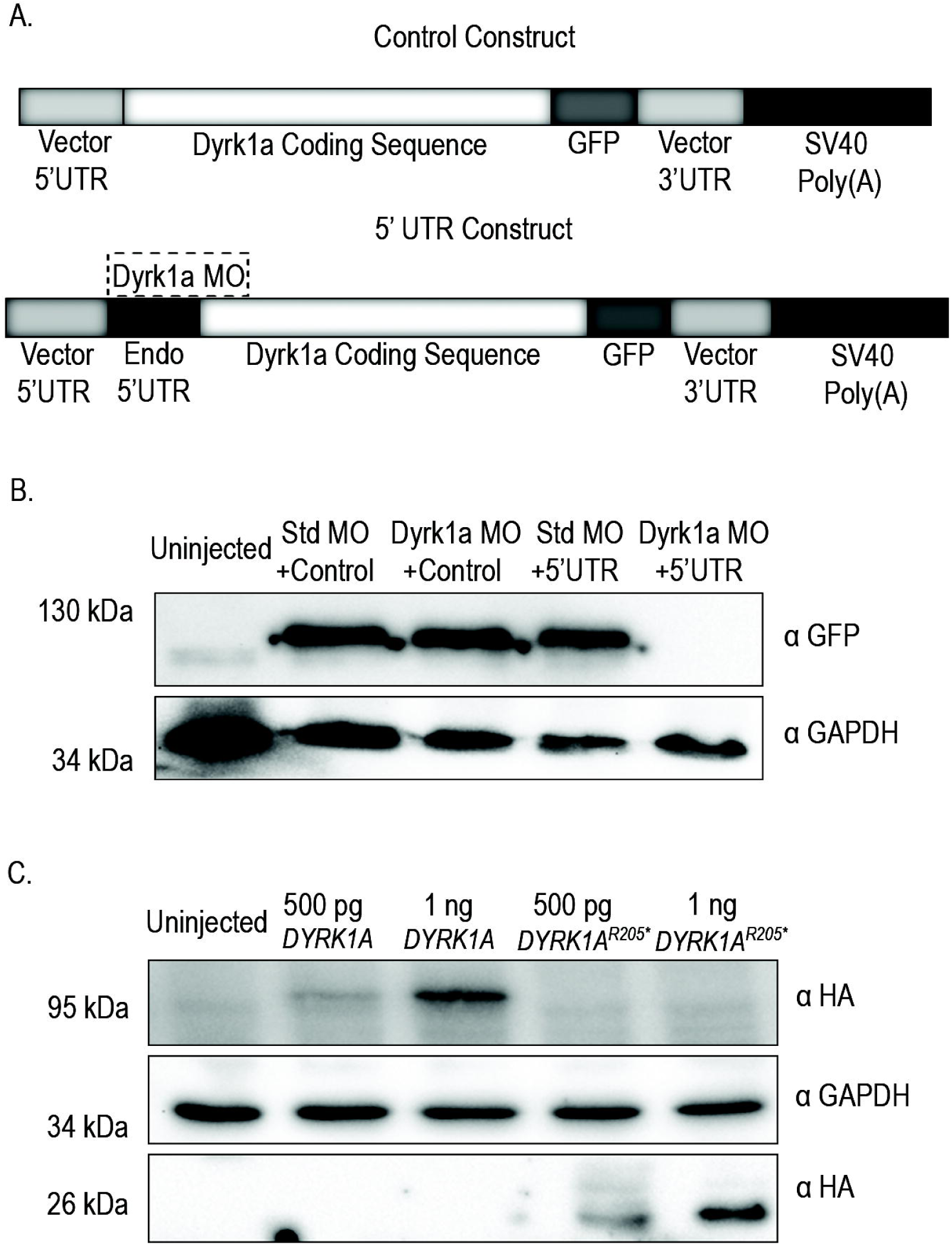
The Dyrk1a MO correctly targets *Xenopus dyrk1a* RNA and both wildtype and truncated human DYRK1A protein are expressed in neurula stage *Xenopus* embryos. Schematic demonstrates that the 5’ UTR construct (A, bottom) contains part of *dyrk1a*’s endogenous 5’ UTR that is recognized by the Dyrk1a MO (shown in dashed box), while the control construct (A, top) does not. Single-cell embryos were injected with 40 ng of either Dyrk1a MO or Standard MO (Std MO) and co-injected with 1 ng RNA (either *Xenopus dyrk1a* control, or *Xenopus dyrk1a* + 5’ UTR) (B). Western blot demonstrates complete reduction of GFP protein levels, indicative of loss of exogenous Dyrk1a, in neurula stage embryos injected with Dyrk1a MO and *dyrk1a +* 5’ UTR RNA (lane 5) compared to the Standard MO and *dyrk1a +* 5’ UTR (lane 4) and Standard MO or Dyrk1a MO and control (lane 2, lane 3) (B). GAPDH was used as a loading control. Wildtype human cDNA was inserted into pCS2-HA vector. *DYRK1A*^*R205\**^ truncated mutation was generated from the wildtype human pCS2-HA-DYRK1A vector and inserted into pCS2-HA vector. Western blot demonstrates HA protein is present in both wildtype (lanes 2 and 3) and truncated *DYRK1A*^*R205*\*^ (lanes 4 and 5) lanes, demonstrating *Xenopus* neurula stage embryos can successfully translate human *DYRK1A* RNA (C). GAPDH was used as a loading control.

**Figure 3.**
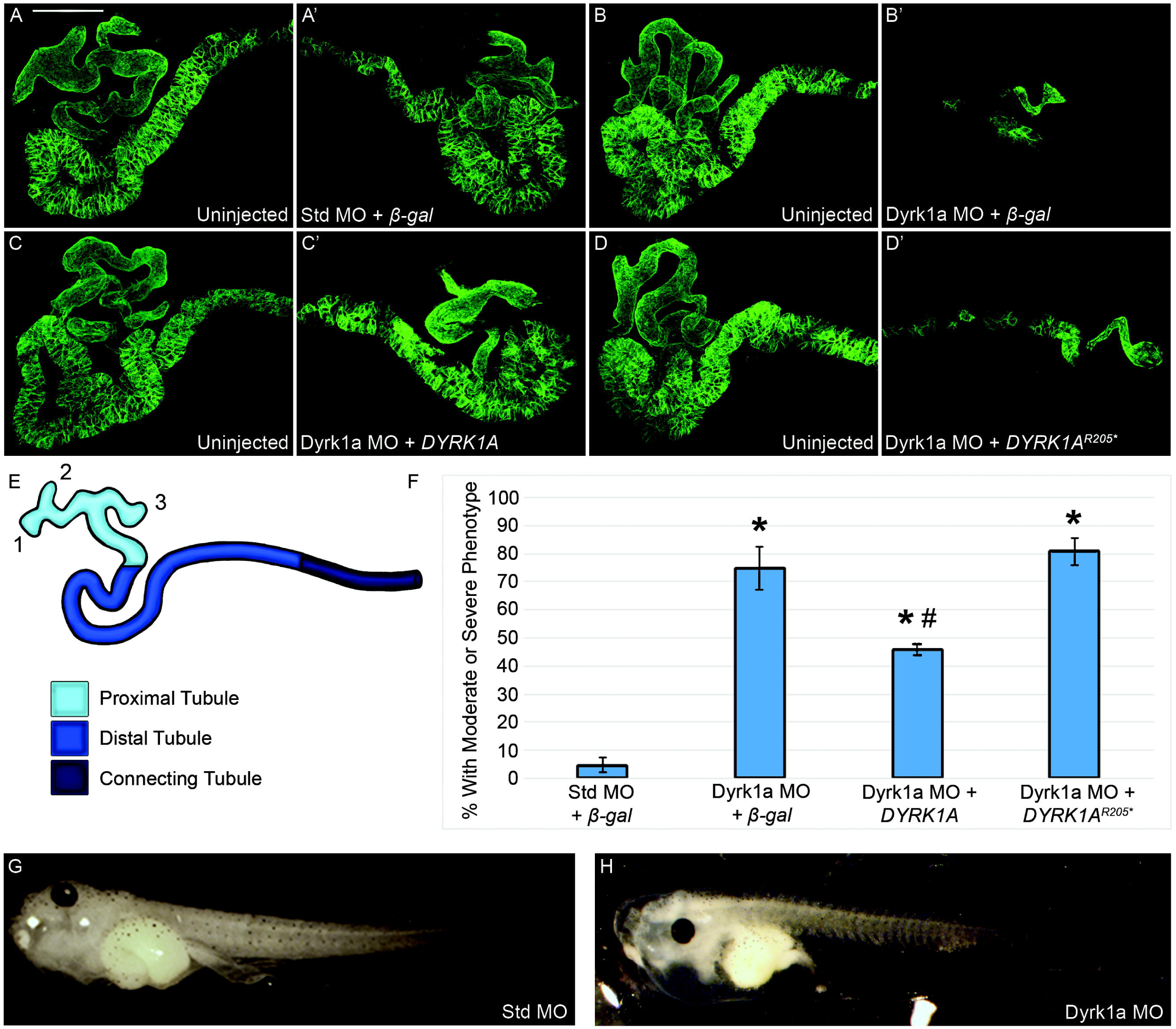
Loss of Dyrk1a affects kidney development in *Xenopus laevis.* Embryos were unilaterally injected at the 8-cell stage with 10ng of Dyrk1a MO or Standard MO (Std MO) along with 50 pg *β-gal*, wildtype or truncated *DYRK1A*^*R205\**^ RNA. Stage 40 tadpoles were stained with kidney antibodies 3G8, which label the proximal tubules and 4A6, which labels the distal and connecting tubules (A-D’). Letters with apostrophes (A’-D’) represent the injected side, whereas letters without apostrophes (A-D) represent the uninjected side. Knockdown with a translation-blocking Dyrk1a MO disrupts kidney development (B’) which can be partially rescued by co-injecting with wildtype human *DYRK1A* RNA (C’) but not truncated *DYRK1A*^*R205\**^ RNA (D’). Co-injection of a Standard MO and β-gal (A’) serve as a negative control. The schematic represents a stereotypical *Xenopus* embryonic kidney (E). The percent of tadpoles missing more than one branching tubule (E numbered 1-3) from the proximal tubule region in light blue (considered a moderate or severe phenotype when two or more parts are missing and is fully characterized in Figure S2) was measured among all four experimental groups (F). Scale bar represents 100 µm. Error bars represent standard error of experiments that were repeated three or more times. The total number of *Xenopus* embryos are as follows: Control: 91, Dyrk1a MO + *β-gal:* 45, Dyrk1a MO + *DYRK1A:* 83, Dyrk1a MO + *DYRK1A*^*R205\**^: 64. * (asterisk) = p<0.0001 compared to Standard MO + *β-gal*. # (pound sign) = p<0.005 compared to Dyrk1a MO + *β-gal*. As expected, no statistical difference was seen between Dyrk1a MO + *β-gal* and Dyrk1a MO *+ DYRK1A*^*R205\**^. Statistical significance was established using a two-tailed T-test. For edema assays embryos were injected at the 4-cell stage in both ventral cells to target both kidneys while avoiding the dorsal cells fated to become the heart and liver, which can also lead to edema. Embryos injected with the Standard MO did not develop edema (G) while embryos injected with the Dyrk1a MO did (H). The total number of *Xenopus* embryos are as follows: Dyrk1a MO: 65 total, 29 of which had edema, Standard MO: 60 total, 0 of which had edema.

### A mutation identified in *DYRK1A*-related ID syndrome fails to rescue Dyrk1a loss-of-function in *Xenopus*

To assess if patient *DYRK1A* mutations lead to pronephric anomalies as they do in *Xenopus*, rescue experiments were carried out upon MO-mediated Dyrk1a knockdown in *Xenopus.* To express human *DYRK1A* in *Xenopus*, constructs with HA tags were generated containing wildtype human *DYRK1A* RNA and truncated human *DYRK1A*^*205\**^ RNA. Western blot analysis with an HA antibody was used to confirm that these human *DYRK1A* constructs could be successfully expressed in *Xenopus* (Figure 2C). The anomalies were partially rescued by coinjecting wildtype human *DYRK1A* RNA (Figure 3C’). However, the patient-identified truncation *DYRK1A*^*205\**^ RNA failed to rescue these anomalies (Figure 3D’). Examples of how embryos are scored can be found in Figure S2. To assess if Dyrk1a affects kidney function a functional assay was performed evaluating edema formation. Edema can be caused by a disruption in the kidneys’ ability to excrete excess fluid, but it can also be caused by heart or liver failure. Both the heart and liver arise from dorsal cells in X*enopus* (Xenbase.org).

To prevent knockdown in these tissues, embryos were injected with Dyrk1a MO or Standard MO in both ventral cells at the four cell stage to affect both kidneys. Embryos injected with the Dyrk1a MO suffered from edema, characterized by swelling in the chest cavity due to fluid retention (Figure 3H) while embryos injected with the Standard MO did not (Figure 3G). This technique establishes that loss of *dyrk1a* affects kidney function in *Xenopus*. Taken together, these data support a role for DYRK1A in pronephric development and strongly suggest that the *DYRK1A*^*R205\**^ mutation is responsible for the kidney anomaly observed in this patient.

## DISCUSSION

Recent discoveries demonstrate that *de novo* pathogenic variants in *DYRK1A* cause a syndromic form of ID [OMIM: 614104]. The findings in the current study indicate that CAKUT/GD should be included as features associated with this syndrome.

CAKUT consist of a heterogeneous clinical spectrum, and how CAKUT arise is largely unknown. Thus, it is important to identify all genes and causal variants involved.^31^ Strong genetic causality of monogenic disease only accounts for 14% of CAKUT cases,^32^ and polygenic causes are speculated to occur but are largely unknown.^33^ Next-generation sequencing, specifically ES, has improved the discovery of novel causative genes that are important in GU development.^34,35,36^ Here, we report on a novel genetic contribution of DYRK1A to CAKUT/GD. We identified 17 unique variants in *DYRK1A* from clinical ES in 19 unrelated individuals. As summarized in Table 1, microcephaly, ID, DD, and seizure are some of the more common features of this syndrome.

Eleven out of fifteen (73% of those with available data) individuals in this study (Table 2) have CAKUT/GD, with 33% having renal anomalies, in addition to other organ involvement. While renal anomalies have been reported previously,^20^ broadly based phenotyping for CAKUT was not performed in previous studies.

Seven of the *DYRK1A* variants identified in our study were novel. Per inclusion criteria, most of *DYRK1A* variants were *de novo* and included truncating variants. This suggests a loss-of-function mechanism for variants causing this syndrome which further supports the findings of another group who proposed reduced kinase function as the cause of this syndrome.^22^

In addition, we studied the role of *dyrk1a* in kidney development in *Xenopus* and found that *dyrk1a* knockdown results in abnormal tubules or complete loss of the kidney. This phenotype could be partially rescued by human *DYRK1A* RNA. However, a nonsense mutation failed to rescue the phenotype, indicating that loss-of-function variants in this gene are, in fact, causative for the observed phenotype in some patients. Furthermore, Dyrk1a MO was injected into two cells to affect both kidneys which resulted in edema suggesting that Dyrk1a is important for both kidney development and function. Although *Xenopus* is an established model to study kidney development, it has not been commonly used to study GD. However, our findings related to pronephric development are likely relevant to the mammalian GU tract, given the dependence of the formation of the male and female urogenital tract upon the nephric duct and the pronephros.

One limitation of this study is that, we were not able to obtain GU information from all patients in this study. Future plans for research include identification and studying the phenotype and underlying variants in a larger number of affected families. Furthermore, signaling pathways involved in *DYRK1A*-related CAKUT/GD should be investigated.

In summary, the phenotype of *DYRK1A*-related ID syndrome is expanded to include CAKUT/GD. Based on our data, we empirically recommend that individuals with DYRK1A syndrome undergo a renal ultrasound and a thorough genital physical exam.

## ACKNOWLEDGEMENTS

We wish to express our sincere gratitude to the patients and their families for their participation in this study. The human studies were supported in part by K12 DK0083014 Multidisciplinary K12 Urologic Research Career Development Program, R01DK078121 from the National Institute of Diabetes and Digestive and Kidney Diseases awarded to D.J.L., startup funding from the Department of Pediatrics (Renal Section to M.R.B.), and the Wood family foundation. We appreciate all the efforts by BG diagnostic laboratory faculty and staff. *Xenopus* studies were supported by National Institute of Diabetes and Digestive and Kidney Diseases grants (K01DK092320 and R03DK118771 to R.K.M.) and startup funding from the Department of Pediatrics, Pediatric Research Center at the McGovern Medical School (to R.K.M.). We thank the instructors and teaching assistants of the 2017 Cold Spring Harbor Laboratory *Xenopus* Course, in particular K.J. Liu and M.K. Khokha. We are grateful to the members of the laboratories of R.K. Miller and P.D. McCrea, as well as to M. Kloc, for their helpful suggestions and advice throughout this project. In particular, we thank H. Ji and P.D. McCrea for valuable constructs and M. Corkins for technical assistance with cloning. J.C. Whitney and T.H. Gomez who took care of the animals, even during hurricane Harvey. We are grateful to the UTHealth Office of the Executive Vice President and Chief Academic Officer and the Department of Pediatrics Microscopy Core for funding the Zeiss LSM800 confocal microscope used in this work. Research reported in this publication was partially supported by the National Institute of Neurological Disorders and Stroke (NINDS) under award number K08NS092898 and Jordan’s Guardian Angels (to G.M.).

## AUTHOR CONTRIBUTIONS

A.T.M.B. cloned *Xenopus* and human *DYRK1A* constructs, performed microinjections for knockdown and rescue experiments, analyzed embryo phenotypes and helped write the manuscript. N.B. contacted referring physicians, collected data, coordinated the study and helped write the manuscript. V.C.M. reviewed clinical data and helped write the manuscript. J.A.R. queried clinical ES data at BG and contacted referring physicians. M.N.B. helped query of ES data. Y.Y. and P.L. interpreted the ES data. Referring physicians include S.M.K., M.R.D., L.H., I.K., D.R.B., P.G.W., L.A., A.M.S.M.A., N.G.O., H.T.C., G.M.M., A.E.S., M.K.K., and F.S. C.E. supervised interpretation of ES data. M.C. B. and D.J.L. supervised research conduction. R.K.M. conceived of the *Xenopus-*related aspect of the project, supported the design of the related experiments, and helped write the manuscript. M.R.B. conceived of the research study, directed research conduction, and helped write the manuscript. All authors critically read and edited the manuscript.

## Figure legend

**Supplementary Figure 1. Human and *Xenopus laevis* protein alignment**. Human *DYRK1A* mRNA transcript variant 1 and *Xenopus laevis dyrk1a* short homeolog mRNA were translated and aligned using Clustal Omega. The overall identity between the protein sequences is 91.3% while the kinase domain has 97.5% identity demonstrating *DYRK1A* is highly conserved. Grey shading represents the kinase domain of *DYRK1A*. Symbols reflect Clustal Omega’s analysis of the residues and examples of specific residue groups that reflect each symbol can be found at their website https://www.ebi.ac.uk/Tools/msa/clustalo/. An * (asterisk) indicates positions with a fully conserved residue. A: (colon) indicates conservation between groups of strongly similar properties. A. (period) indicates conservation between groups of weakly similar properties. DYRK1A encodes a protein of 763 or 754 amino acid residues which results from alternative splicing with the longer isoform representing the canonical sequence shown above. *Xenopus* is missing residues 70-78 which are the same residues the short isoform of human DYRK1A (754 amino acids) is missing. This suggests that *Xenopus* most likely expresses the short isoform of DYRK1A.

**Supplementary Figure 2. Scoring system for *Xenopus* embryonic kidneys**. Kidneys were scored at Nieuwkoop and Faber stage 40.^11^ A normal kidney consists of three branches in the proximal region with a “trunk” connecting to the convoluted looping of the early distal region, and strong immunostaining of the late distal region (A). A weak phenotype consists of loss of one of the branches in the proximal region and possibly less early distal looping and/or partial loss of late distal immunostaining (B). A moderate phenotype consists of loss of two of the branches in the proximal region with moderate loss of early distal looping and possible loss of late distal immunostaining (C). A severe phenotype consists of complete loss of proximal tubules with severe or complete loss of early distal looping and the late distal region (D).

