## Supplementary material for "*DYRK1A*-related intellectual disability: a syndrome associated with congenital anomalies of the kidney and urinary tract": Supp Fig. 1

```
Human DYRK1A        MHTGGETSACKPSSVRLAPSFSFHAAGLQMAGQMPHSH-QYSDRRQPNISDQQVSALSYS    59
Xenopus dyrk1a      MHTGGETSACKPSSVRLAPSFSFHAAGLQMAGQMSHSHQQYSDRHQQNLNDQQASALPYN    60
                    ********************************** *** *****:* *:.***.*** *.

Human DYRK1A        DQIQQPLTNQVMPDIVMLQRRMPQTFRDPATAPLRKLSVDLIKTYKHINEVYYAKKKRRH    119
Xenopus dyrk1a      DQTPQPLPNQ---------RRMPQTFRDPATAPLRKLSVDLIKTYKHINEVYYAKKKRRH    111
                    **  *** **         *****************************************

Human DYRK1A        QQGQGDDSSHKKERKVYNDGYDDDNYDYIVKNGEKWMDRYEIDSLIGKGSFGQVVKAYDR    179
Xenopus dyrk1a      QQGQGDDSSHKKERKVYNDGYDDDNYDYIVKNGEKWMDRYEIDSLIGKGSFGQVVKAYDR    171
                    ************************************************************

Human DYRK1A        VEQEWVAIKIIKNKKAFLNQAQIEVRLLELMNKHDTEMKYYIVHLKRHFMFRNHLCLVFE    239
Xenopus dyrk1a      VEQEWVAIKIIKNKKAFLNQAQIEVRLLELMNKHDTEMKYYIVHLKRHFMFRNHLCLVFE    231
                    ************************************************************

Human DYRK1A        MLSYNLYDLLRNTNFRGVSLNLTRKFAQQMCTALLFLATPELSIIHCDLKPENILLCNPK    299
Xenopus dyrk1a      MLSYNLYDLLRNTNFRGVSLNLTRKFAQQMCTALLFLATPELSIIHCDLKPENILLCNPK    291
                    ************************************************************

Human DYRK1A        RSAIKIVDFGSSCQLGQRIYQYIQSRFYRSPEVLLGMPYDLAIDMWSLGCILVEMHTGEP    359
Xenopus dyrk1a      RSAIKIVDFGSSCQLGQRIYQYIQSRFYRSPEVLLGTPYDLAIDMWSLGCILVEMHTGEP    351
                    ************************************ ***********************

Human DYRK1A        LFSGANEVDQMNKIVEVLGIPPAHILDQAPKARKFFEKLPDGTWNLKKTKDGKREYKPPG    419
Xenopus dyrk1a      LFSGANEVDQMSKIVEVLGIPPAHILDQAPKARKFFEKMPEGTWNLKKTKDGKKEYKPPG    411
                    ***********.**************************:*:************:******

Human DYRK1A        TRKLHNILGVETGGPGGRRAGESGHTVADYLKFKDLILRMLDYDPKTRIQPYYALQHSFF    479
Xenopus dyrk1a      TRKLHNILGVENGGPGGRRAGESGHTVADYLKFKDVILRMLDYDAKTRIQPYYALQHSFF    471
                    ***********.***********************:******** ***************

Human DYRK1A        KKTADEGTNTSNSVSTSPAMEQSQSSGTTSSTSSSSGGSSGTSNSGRARSDPTHQHRHSG    539
Xenopus dyrk1a      KKTADEGTNTSNSVSTSPAMEQSQSSGTTSSTSSSSGGSSGTSNSGRARSDPTHQHRHSG    531
                    ************************************************************

Human DYRK1A        GHFTAAVQAMDCETHSPQVRQQFPAPLGWSGTEAPTQVTVETHPVQETTFHVAPQQNALH    599
Xenopus dyrk1a      GHFTTAV-AMDCETHSPQVRQQFP--PGWTVPEAPTQVTIETHPVQETTFHVPPSQKNVP    588
                    ****:** ****************   **:  *******:************ *.*: :

Human DYRK1A        HHHGNSSHHHHHHHHHHHHHGQQALGNRTRPRVYNSPTNSSSTQDSMEVGHSHHSMTSLS    659
Xenopus dyrk1a      HHHGNGSHH--HHHHHHHHHGQHILSNRTRTRIYNSPSTSSSTQDSMDIGNSHHSMTSLS    646
                    *****.***  ***********: *.**** *:****:.********::*:*********

Human DYRK1A        SSTTSSSTSSSSTGNQGNQAYQNRPVAANTLDFGQNGAMDVNLTVYSNPRQETGIAGHPT    719
Xenopus dyrk1a      SSTTSSSTSSSSTGNQGNQAYQNRPVAANTLDFGQNGTLDVNLTVYSNPRQETGITGHPD    706
                    *************************************::****************:***

Human DYRK1A        YQFSANTGPAHYMTEGHLTMRQGADREESPMTGVCVQQSPVASS                    763
Xenopus dyrk1a      YQYSANTGPGHYVTEGQLTMRQGIDREDSPMTGVCVQQSPVASS                    750
                    **:******.**:***:****** ***:****************
```
