## Supplementary figures and images for "*DYRK1A*-related intellectual disability: a syndrome associated with congenital anomalies of the kidney and urinary tract"

### Supp Fig. 2

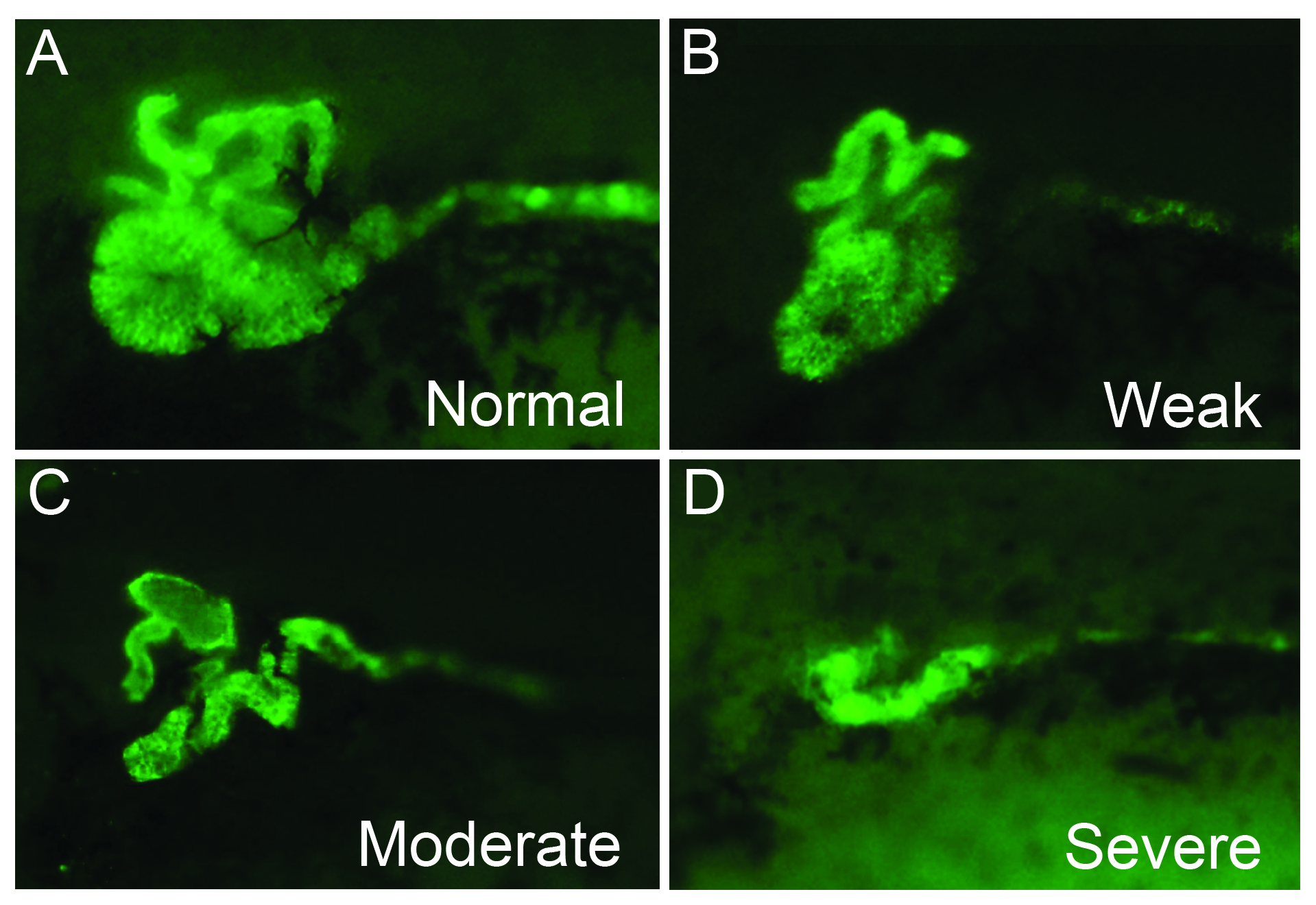
